# Potential G-quadruplex forming sequences and *N*^6^-methyladenosine colocalize at human pre-mRNA intron splice sites

**DOI:** 10.1101/2020.02.07.939116

**Authors:** Manuel Jara-Espejo, Aaron M. Fleming, Cynthia J. Burrows

## Abstract

Using bioinformatic analysis of published data, we identify in human mRNA that potential G-quadruplex forming sequences (PQSs) colocalize with the epitranscriptomic modifications *N*^*6*^-methyladenosine (m^6^A), pseudouridine (Ψ), and inosine (I). A deeper analysis of the colocalized m^6^A and PQSs found them intronic in pre-mRNA near 5′ and 3′ splice sites. The loop lengths and sequence context of the m^6^A-bearing PQSs found short loops most commonly comprised of A nucleotides. This observation is consistent with literature reports of intronic m^6^A found in SAG (S = C or G) consensus motifs that are also recognized by splicing factors. The localization of m^6^A and PQSs in pre-mRNA at intron splice junctions suggests that these features could be involved in alternative mRNA splicing. A similar analysis for PQSs around sites of Ψ installation or A-to-I editing in mRNA also found a colocalization; however, the frequency was less than that observed with m^6^A.

**TOC Graphic:** 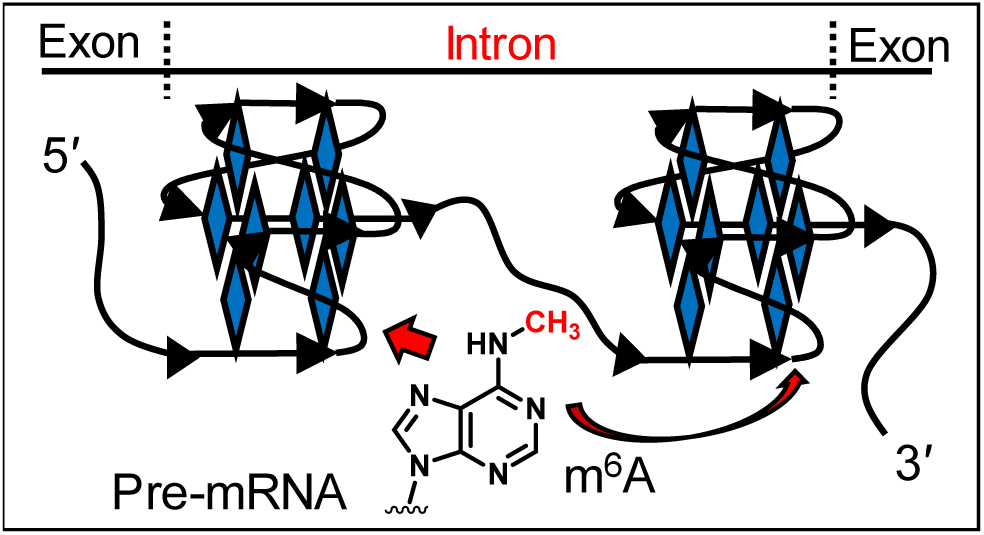

---

Interest in the fate of nascent mRNA has focused on sequence and structure to either attract or repel protein readers and drive sites of writing chemical modification.^1-5^ Maturation of mRNA involves installation of a 5′ cap, addition of a 3′-polyadenosine tail, installation of epitranscriptomic modifications, and intron excision.^6^ Processing of the 5′ and 3′ ends of mRNA has long been studied.^7,8^ In contrast, our understanding of writing epitranscriptomic modifications and alternative mRNA splicing are rapidly advancing as a result of next-generation sequencing (NGS) and expansion of bioinformatic tools.^1-4^ The best studied epitranscriptomic modification in mRNA is *N*^*6*^-methyladenosine (m^6^A; Figure 1A), in which deposition of the methyl group on A nucleotides occurs in specific sequence motifs; however, not all such sequences are modified.^1-5^ This has led to the suggestion that RNA secondary structure plays a key role in selection of m^6^A modified sites.^9-11^ Additional epitranscriptomic modifications include pseudouridine (Ψ) and A-to-I editing yielding inosine (I; Figure 1A), in which similar lingering questions regarding the sites selected for modification have been asked.^9^

**Figure 1.**
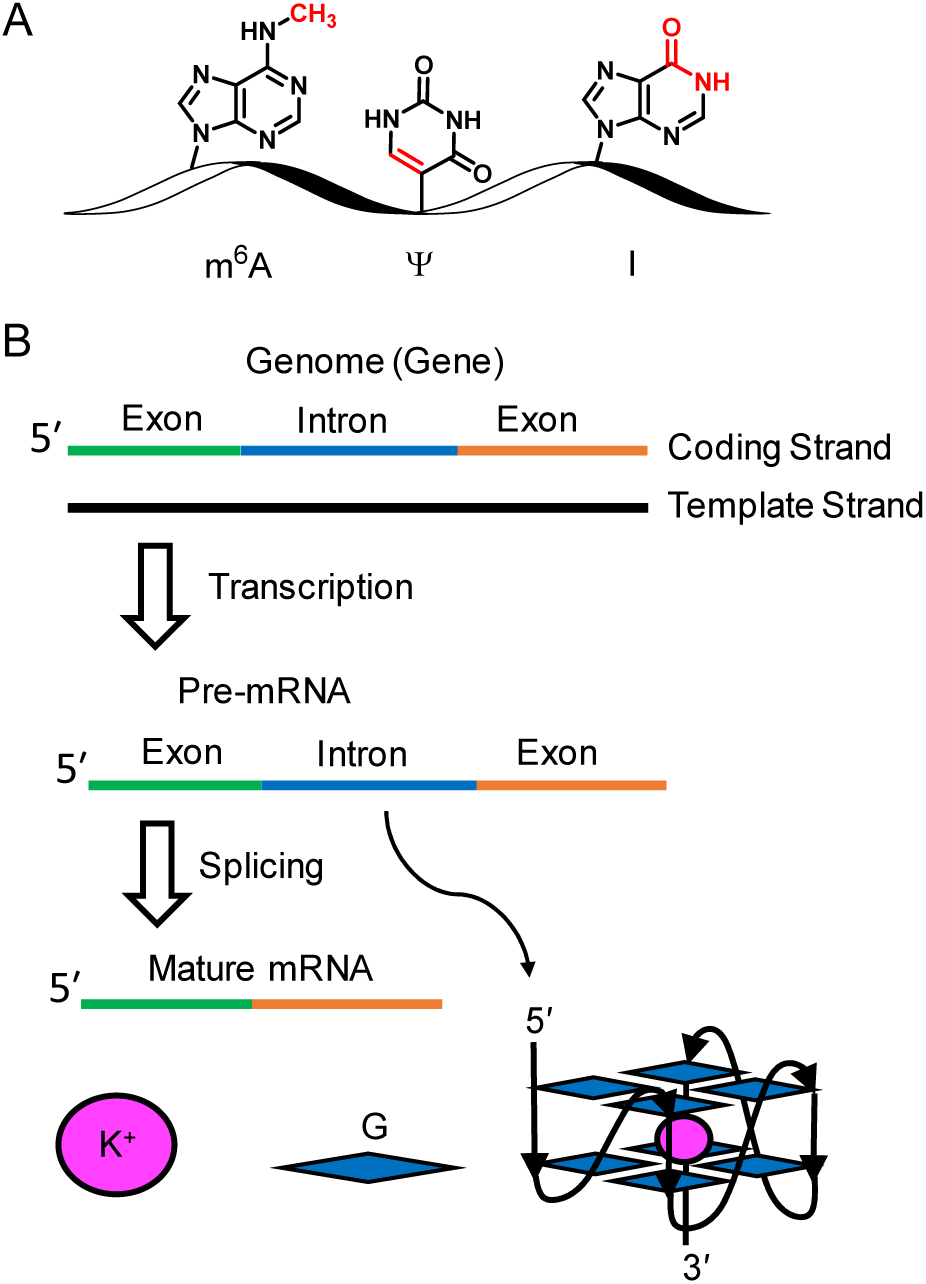
(A) The modifications m^6^A, Ψ, and I are epitranscriptomic. (B) Maturation of mRNA involves intron splicing; potential G-quadruplex sequences are found in introns.

Alternative splicing of nascent or pre-mRNA to yield mature mRNA is a highly regulated eukaryotic process resulting in a single gene being able to code for multiple proteins.^12^ This diversification occurs by inclusion or exclusion of particular exons in the final processed mRNA. In humans, ∼95% of multi-exonic genes are alternatively spliced enabling the ∼20,000 protein-coding genes to direct synthesis of a much greater diversity of proteins. Alternative splicing of mRNA is cell type specific and changes with oxidative stress or disease.^12,13^ The mRNA features and epitranscriptomic components that drive when and to what extent alternative splicing occurs in particular cell types have been topics of recent studies,^12,13^ and future work is needed to better understand the RNA structural details guiding the process.

Genomes code for all the information found in RNA (Figure 1B), and a fascinating finding in the human genome is enrichment of potential G-quadruplex forming sequences (PQSs) in promoters, 5′-untranslated regions, and introns near splice sites.^14^ G-Quadruplexes (G4s) are non-canonical folds in nucleic acids that have sequences comprised of four or more runs of G in close proximity providing the opportunity for the sequence to fold around intracellular K^+^ ions (Figure 1B).^15^ In DNA G4s, stable folds each have G-runs of three or more Gs, while in RNA G4s (rG4s), two Gs per run can adopt stable folds. Those PQSs in the non-template strand (i.e., coding strand) of a gene will also be present in the RNA transcript (Figure 1B). The role of PQSs and their folding to rG4s to guide alternative splicing of specific mRNA, such as the p53 mRNA, has been noted.^16,17^ Moreover, sequencing m^6^A in nascent mRNA has found enrichment around splice junctions, in addition to those near the start and stop codons.^18,19^ Biochemical studies suggest m^6^A is written on nascent mRNA as it is synthesized, and the m^6^As in introns are lost in the mature mRNA.^18^ Herein, using bioinformatic analysis of published data, we identify that PQSs in pre-mRNA colocalize with sites of the epitranscriptomic modification m^6^A in introns, suggesting a possible synergy in deposition of m^6^A on pre-mRNA around PQSs. A similar analysis for PQSs around sites of Ψ installation or A-to-I editing in mRNA was conducted to also find a colocalization; however, the colocalization of these modifications with PQSs was less abundant than observed with m^6^A.

The publicly available sources for the RNA modification maps in human mRNA used in the present study are summarized in Table S1.^9,18-20^ The peak summits for each epitranscriptomic modification were identified, and we then selected a window of sequence space ± 30 nucleotides flanking the summit to inspect computationally for PQSs. The quadruplex-forming G-rich sequence (QGRS) mapper algorithm was used to look for the sequence pattern 5′-G_x_ L G_x_ L G_x_ L G_x_-3′ where x ≥ 2 nucleotides and L represents the loops with a total length = 22 nucleotides.^21^ The first dataset inspected had sequenced m^6^A in mature mRNA collected from HeLa cells using the m^6^A-CLIP sequencing protocol.^19^ From these data in mature HeLa mRNA, 17% (7,838 out of 46,355) of the m^6^A enriched regions were found to also contain a PQS (Figure 2A). Guided by the knowledge that PQSs are enriched in human introns,^22^ inspection of the chromatin-associated RNA (i.e., pre-mRNA) in HeLa cells from the same report were interrogated.^19^ A similar colocalization of PQSs and m^6^A was observed in the pre-mRNA as found in the mature mRNA sequencing datasets (21% vs. 17%; Figure 2A). Next, there exists another available dataset for m^6^A in pre-mRNA from HEK293 cells that was sequenced using transient *N*^*6*^-methyladenosine transcriptome sequencing (TNT-seq).^18^ In 40% (23,372 out of 58,311) of the enriched m^6^A sites found in the HEK293 pre-mRNA, a PQS was also located in the same region. The analyses of literature data, particularly the studies in HEK293 cells, suggest a high incidence of m^6^A enriched sites occurring in mRNA regions that have a PQS likely to adopt a rG4 fold.

**Figure 2.**
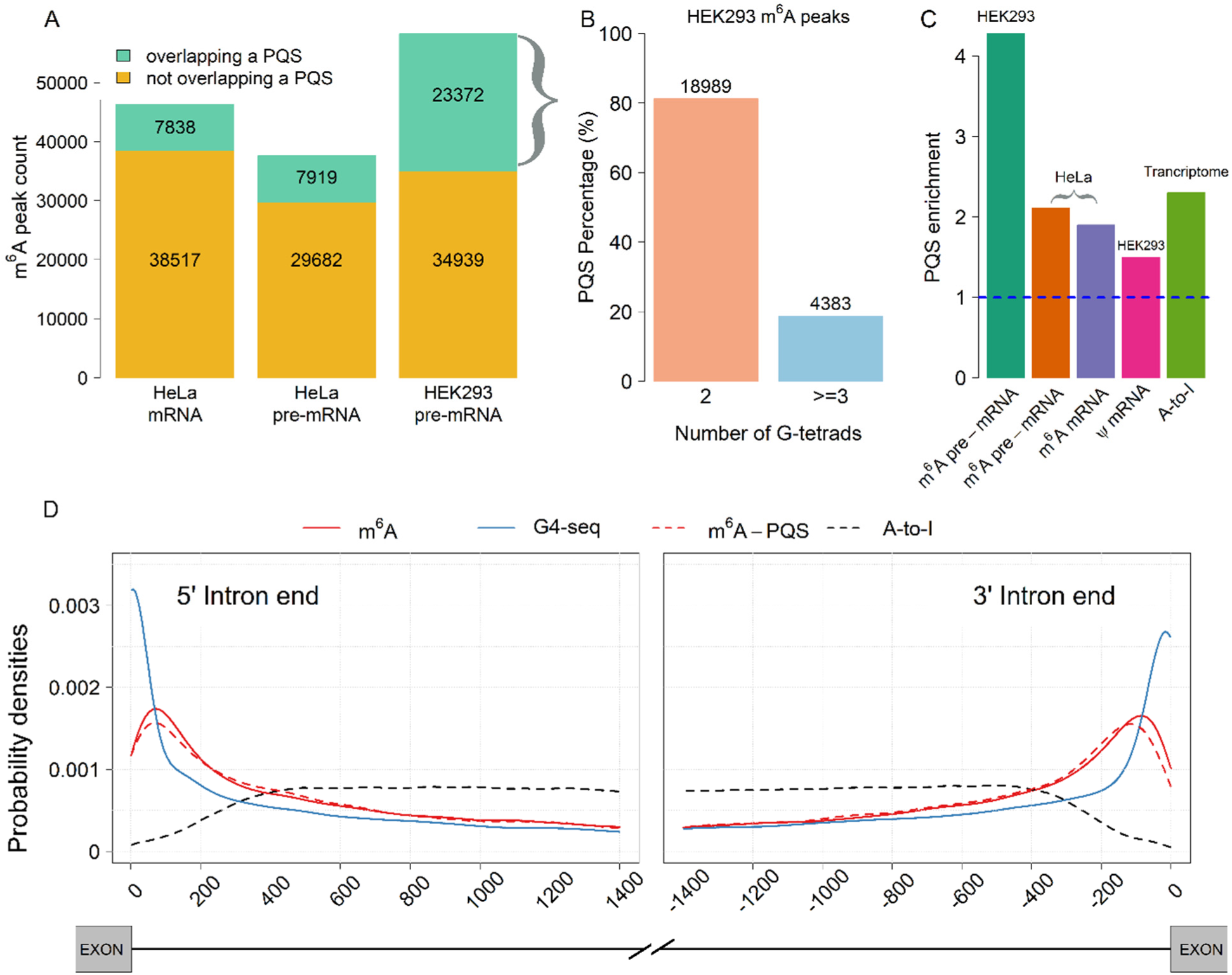
Inspection of human mRNA sequenced for m^6^A, Ψ, and A-to-I editing sites to determine whether a PQS resides to the same location. (A) Bar plot showing the number of m^6^A enriched sites in HeLa and HEK293 mRNA that also have a PQS overlapping with the site of modification. (B) Classification of the PQSs found in the HEK293 pre-mRNA on the basis of the number of G-tetrads that can form from the sequence. (C) Plot of PQS enrichment (observed/expected) found in the epitranscriptomic modification sites. (D) Intron map illustrating the m^6^A enriched sites found in HEK293 pre-mRNA sequenced by TNT-seq,^18^ the position of the PQSs found in the sequencing data, and comparison to the position of G4s found in the human genome via G4-seq.^22^ This plot also provides a map of the A-to-I editing sites in human introns.^20^

In the next step of the analysis, the PQSs found in the HEK293 pre-mRNA sites of m^6^A enrichment were classified on the basis of the number of G-tetrads that would occur in the G4 fold. The PQS population divided out to 81% having two G-tetrads and 19% having three or more G-tetrads (Figure 2B). A similar breakdown of the two versus three G-tetrad G4s was found in the PQSs identified in the HeLa pre-mRNA and HEK293 mature mRNA m^6^A and Ψ datasets analyzed (Figure S1). In RNA, stable rG4 folds have been found with only two G-tetrads, which is in contrast to DNA that generally requires at least three G-tetrads to adopt a stable fold.^10,15^ The greater stability in rG4s results from the 2′-OH providing an additional hydrogen bond that is not possible in DNA.^15^ Prior studies have shown that two-tetrad rG4s provide a quasi-stable fold that can be harnessed as a switch to impact the fate of RNA in cells.^23^ The finding of m^6^A and PQS colocation in human mRNA, particularly introns of pre-mRNA, suggests that the rG4 secondary structure and epitranscriptomic m^6^A may be synergistic; this observation is consistent with our previous report on a similar finding in viral genomic RNA.^10^ Whether the G4 fold is the signal for writing m^6^A in the mRNA or the presence of m^6^A impacts the G4 fold is not known.^10^

The statistical significance for enrichment of PQSs in the regions of m^6^A enrichment in the human mRNA was compared to randomized, shuffled sequences (Figure 2C). Comparison of the 23,372 PQSs identified in the HEK293 pre-mRNA to the 5,461 PQSs expected by randomized shuffling found a 4.3-fold enrichment in PQSs; this finding is significant on the basis of Fisher’s exact test (*P* < 2.2 e^-16^; Figure 2C and Table S2). In the HeLa mRNA analyzed for m^6^A and PQS colocalization, there was a significant 2.1-fold enrichment found in the pre-mRNA (*P* < 2.2 e^-16^; Figure 2C and Table S2) and a significant 1.9-fold enrichment found in the mature mRNA (*P* < 2.2 e^-16^; Figure 2C and Table S2). Taken together, these results support a favorable colocalization of m^6^A sites and PQSs in human mRNA.

Analysis of epitranscriptomic maps for Ψ and A-to-I editing in human mRNA for colocalization with PQSs was conducted; however, a Ψ map in pre-mRNA is not yet available. Therefore, the PQS analysis was conducted on mature mRNA from HEK293 cells sequenced for Ψ (Table S1).^9,20^ In the Ψ dataset, 16% of the modified sites (325 out of 2,058) also contained a PQS. Surprisingly, this number of PQSs represents a significant 1.5-fold enrichment of these G-rich sequences (*P* < 8.8 e^-08^; Figure 2C and Table S2). In the A-to-I analysis, 22% (1,004,026 out of 4,668,508) colocalized with a PQS (Figure 2C) that was a significant 2.3-fold enrichment (*P* < 2.2 e^-16^; Table S2). The G-tetrad count for PQSs colocalized with Ψ or A-to-I editing sites was also ∼80% two G-tetrad G4s and ∼20% three or more G-tetrad G4s (Figure S1); these values are very similar to those found in the m^6^A PQS colocalization data.

In the HEK293 pre-mRNA m^6^A data reported by Louloupi, et al.,^18^ a high incidence of m^6^A residing in intronic regions was observed. Next, a focused inspection of PQSs in the intron regions was conducted. As expected, the m^6^A data (Figure 2D solid red) and PQS (Figure 2D dashed red) data tracked with each other and were favorably enriched on the intronic side of both the 5′ and 3′ splice sites. The Balasubramanian laboratory developed G4-seq to find all sequences that could adopt G4s in the human genome.^22^ Our next analysis of the intronic data looked for folded *genomic* G4s, identified experimentally, that track with m^6^A and PQSs in RNA. There is one noteworthy point regarding this comparison—stable DNA G4s are generally at least three G-tetrads, while in the RNA analysis here a preference for two G-tetrad G4s was observed (Figures 2B and S1). Nevertheless, we expected to see at least 20% of the data. With this limitation in mind, the G4-seq data on the coding strand of human introns did indeed show enrichment at both the 5′ and 3′ splice sites (Figure 2D blue line) that also tracked with the m^6^A and PQS profiles. This new observation strengthens the argument of a colocalization of m^6^A and PQSs around intron splice sites.

A closer examination of the data around intron splice sites provided a few interesting observations. Within 200 nt of each splice junction, 8,040 m^6^A-enriched sites on the intronic 5′ splice site representing 3,187 genes and 7,236 m^6^A-enriched sites on the intronic 3′ splice site representing 2,681 genes were found; these numbers represent 47% of total m^6^A intronic sites in the HEK295 pre-mRNA dataset. A similar distribution of m^6^A and PQS colocalization sites was observed indicating a possible close association of these two RNA features near mRNA splicing sites. These observations suggest an exciting opportunity for future studies that address whether rG4s and m^6^A function synergistically to guide mRNA splicing.

The A-to-I editing sites in introns of HEK293 mRNA were plotted alongside the m^6^A, PQS, and G4-seq data that found RNA editing was not observed around splice sites and did not track with PQSs in this region (Figure 2D black dashed line). The A-to-I editing sites appear to be depleted around intron splicing sites. The Ψ dataset from HEK293 cells was conducted on mature mRNA, in which introns are not present, and therefore, no further analysis of the data was conducted.

In mature mRNA, m^6^A mapping studies have suggested a broad consensus motif for **A** methylation in the sequence context DR**A**CH (D = A, G, or U; R = G or A; and H = A, C, or U). In the work by Louloupi, et al., intronic m^6^A was favorably deposited in S**A**G (S = C or G) sequence motifs.^18^ Because the HEK293 pre-mRNA data exhibited the highest colocalization of m^6^A and PQSs, this PQS population was further interrogated to identify favorable G4 loop profiles, with respect to length and sequence. During the bioinformatic PQS inspection, the three loops could have a total length of 22 nucleotides or roughly 7 or fewer nucleotides per loop. The loop analysis was conducted on 14,076 two G-tetrad PQSs and 729 three or more G-tetrad PQSs. In the loop length analysis of two G-tetrad PQSs, there was a slight preference for shorter loop lengths, but there were many longer loop PQSs observed at a high relative frequency (Figure 3A). Additionally, a breakdown of the 1^st^, 2^nd^, or 3^rd^ loops found they all had a similar length profile (Figure 3A). In the PQS population with three or more G-tetrads, one nucleotide loop lengths were less common, while three and four nucleotide loop lengths were most common (Figure 3B). Inspection of the combination of all three loop lengths together in the two G-tetrad PQSs found the 1-1-1 loop length combination to be most common followed by the 4-4-4 loop length combination. In general, as the loop length combination increased or became asymmetric, the number of PQSs observed decreased (Figure 3C). For the three-loop length combination analysis of the three G-tetrad or more PQSs, the most common combination found had 3-3-3 nucleotide loop lengths; this was followed by the longer 7-7-7 and shorter 2-2-2 nucleotide loop lengths (Figure 3D). In the three or more G-tetrad PQS data, the least common were those with asymmetric loop lengths, with the exception to the least common pool being 6-6-6 nucleotide long loops (Figure 3D).

**Figure 3.**
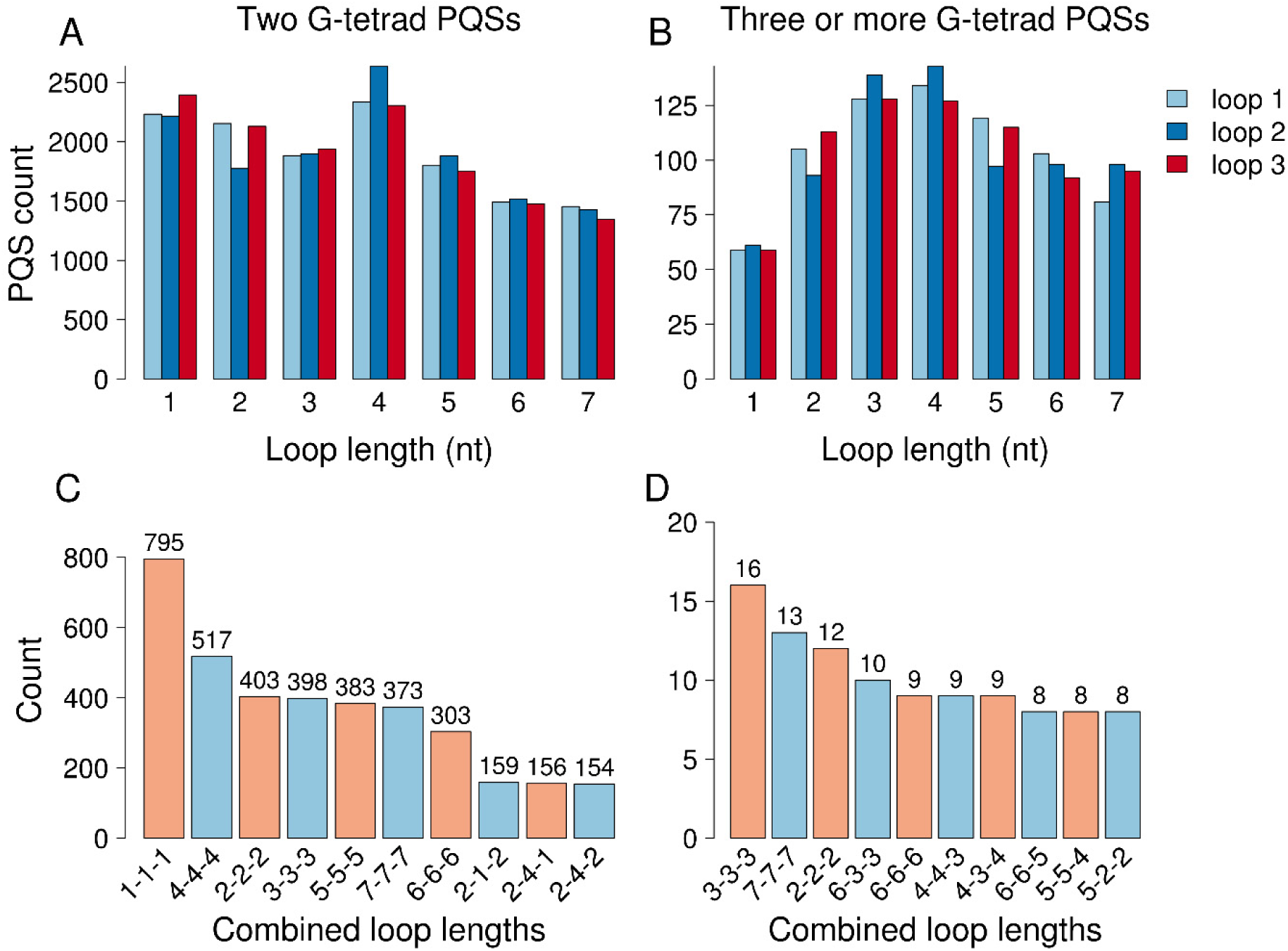
Loop length analysis of the PQSs that colocalized with m^6^A enriched regions in the HEK293 pre-mRNA.^18^ Analysis of individual loop lengths for (A) two G-tetrad PQSs and (B) three or more G-tetrad PQSs. Analysis of loop length combinations for (C) two G-tetrad PQSs and (D) three or more G-tetrad PQSs that identify the most prevalent loop length combinations.

Inspection of the loop sequences of the two G-tetrad PQSs in m^6^A-enriched regions identified a nucleotide preference. In Table 1 a rank ordering of the top five most common loop sequences found in each of the three possible G4 loops is provided. Single-nucleotide loops comprised of an A nucleotide were the most common in all three loops. Because A nucleotides are potential sites of methylation, the observation of single A nucleotides in the two G-tetrad PQSs nicely fits our hypothesis that G4 folds provide a structural motif to guide sites of m^6^A introduction in human mRNA. Furthermore, the high incidence of single A nucleotides is consistent with the work by Louloupi, et al.,^18^ in which the consensus motif S**A**G occurred with greater frequency. This observation leads to future experimental questions: (1) This bioinformatic study suggests a synergy between rG4 folds with sites of m^6^A epitranscriptomic modification. Do rG4 folds function as a structural motif for methylation of the RNA by the METTL3/14 methyltransferase complex? (2) The S**A**G consensus sequence is also recognized by some splicing proteins such as SRSF3.^18^ Do these splicing proteins bind rG4s, and is their binding modulated by the presence of m^6^A in a rG4 loop? (3) Lastly, splicing factors that bind S**A**G sequence motifs were found to be involved in alternative mRNA splicing.^24^ Is there synergy between rG4s and m^6^A to guide alternative mRNA splicing? This final question has been studied by looking at rG4s,^16,17^ but the inclusion of m^6^A may have an additional unknown impact.

**TABLE 1.** PQS loop nucleotide composition for those in m^6^A-enriched regions*

|  | Most to Least Common |  |  |  |  |
| --- | --- | --- | --- | --- | --- |
| Order | 1 | 2 | 3 | 4 | 5 |
| LOOP 1 | A | U | GA | G | AA |
| Count | 1044 | 518 | 502 | 474 | 237 |
| LOOP 2 | A | U | G | GA | C |
| Count | 936 | 657 | 384 | 250 | 239 |
| LOOP 3 | A | U | G | CA | GA |
| Count | 999 | 691 | 459 | 379 | 332 |

\*Data corresponds to HEK293 pre-mRNA m<sup>6</sup>A profile reported by Louloui, et al.<sup>18</sup> See Table S3 for additional data.

The second most common loop sequence observed in the two G-tetrad PQSs was single U nucleotides in all three loops. Many of the top ten most prevalent loop sequences contained A nucleotides within dinucleotide motifs such as 5′-GA, 5′-AA, 5′-CA, and 5′-AG. Interestingly, the 5′-AC dinucleotide that would indicate the DR**A**CH consensus motif within a PQS was not among the top 10 most prevalent loop sequences (Table S3). Further inspection of the 795 PQSs in which single-nucleotide loops were identified also found that homogenous loops of single A nucleotides dominated the distribution with 142 occurrences. Inspection for homogenous single U or C nucleotide loops found 18 and 34 occurrences, respectively, and no PQSs were observed with all-G loops. In summary, two G-tetrad PQSs in m^6^A enriched regions are biased in their nucleotide composition to have A > U > G > C and in their propensity to have the same lengths for all three loops (Figure 3 and Tables 1 and S3). This indicates that homogeneous short loops favor A nucleotides as evidenced by the loop composition analysis, although the sequences with all C or U nucleotide loops will not provide an RNA substrate for writing m^6^A. These sequences may be false positives or the methylated A may reside just beyond the G4; additional analysis of the tail sequences for the small sample of PQSs without an A was not conducted. The sequence composition for the three on more G-tetrad PQSs found colocalized with m^6^A were also analyzed and found to be rich in A nucleotides (Table S4). Additionally, the 5′-AC dinucleotide common to the DR**A**CH consensus motif was not found in the top ten most common loop sequences.

Herein, we explored the structural pattern of human RNA sites harboring m^6^A, Ψ, or A-to-I editing modifications, focusing on the presence of PQSs at these sites. The study revealed that m^6^A-enriched sites colocalize with PQSs, and more interestingly, the greatest colocalization was for intronic sites of m^6^A deposition near splice sites in pre-mRNA. This observation suggests there may be an interplay between, m^6^A, rG4s, and mRNA splicing that could be a component of alternative splicing; future experimental work is needed to address this possibility. Our prior interest in the colocalization of m^6^A and PQSs focused on viral RNA genomes that showed a preference for DR**A**CH motif methylation.^10^ The present work on human pre-mRNA is consistent with the viral RNA analysis; however, the sequence context found for the colocalization in intronic pre-mRNA occurs largely within S**A**G motifs. This difference observed may reflect the writing of m^6^A on pre-mRNA occurs in the nucleus, while m^6^A in viral RNA occurs in the cytosol.^10,18^ Finally, it is important to note that other RNA modifications, such as Ψ and A-to-I editing, also overlapped with PQSs, although at a lower frequency when compared to m^6^A (Figure 2C).

## Associated Content

### Supporting Information

The Supporting Information is available free of charge at https://pubs.acs.org/doi/XXX

Complete methods, Tables for data sources, complete statistical values, and loop sequence composition data.

### Notes

The authors declare no competing financial interests.

## Acknowledgments

This work was funded by a grant from the U.S. National Institute of General Medical Sciences (R01 GM093099). MJE thanks Prof. Sergio Line and the Brazilian Coordination for the Improvement of Higher Education Personnel-CAPES-PRINT 88887.364735/2019-00 for supporting his stay at the University of Utah.

