## Supporting Information for "Potential G-quadruplex forming sequences and *N*^6^-methyladenosine colocalize at human pre-mRNA intron splice sites"

| Item | Page |
|---|---|
| **Complete Methods** | S2 |
| **Table S1**. Details of the literature data used in the analysis | S3 |
| **Table S2.** Fisher exact test results for PQS enrichment analysis | S4 |
| **Table S3.** Two G-tetrad PQS loop nucleotide composition for those in $m^6A$ enriched regions | S5 |
| **Table S4**. Frequency analysis of loop sequences in three or more G-tetrad PQSs in intronic $m^6A$ enriched sites. | S6 |
| **Figure S1**. Frequency of two and three or more G-tetrad PQSs in mature mRNA sequenced for $m^6A$, Ψ, and A-to-I editing. | S7 |
| **References** | S8 |

### Complete Methods

#### RNA modification data sources and analyses

The RNA modification data sources used in the analysis are provided in Table S1. For each RNA modification data set, a 60-nt DNA sequence window centered at the peak summit of enrichment for each modification was selected. The coordinates for the window analyzed were derived from the human hg19 reference genome using the twoBitToFasta tool.[1] The pool of sequences provided the experimental sequence data utilized in the bioinformatic analysis. Additionally, these modification-containing regions were randomly genome-wide positioned (n=10 times) using the bedtools suite.[2]

For the potential G-quadruplex forming sequences (PQS), their identification in the extracted RNA sequences were analyzed using the quadruplex-forming G-rich sequence (QGRS) mapper algorithm.[3] The QGRS mapper uses a motif definition of 5′-$G_{x1}L_1G_{x2}L_2G_{x3}L_3G_{x4}$, where $G_{xn}$ stands for G-run number and $L_n$ represents the loop sequence between each G run. The length of the G-runs was defined as $x \geq 2$ nucleotides and the algorithm allows at most one of the loop lengths to be a length = 0. The same analysis was applied to the shuffled (i.e., randomized) sequences to test for PQS enrichment. Significance for the relative peak enrichment that is defined as the number of PQSs in the experimental versus random-generated sequences was determined using the Fisher test. The coordinates for the genomic G4s found by G4-seq were obtained from GEO accession number GSE63874.[4] The data from this source were analyzed using bedtools running the closest command to identify the nearest G4 to the intron ends.

**Table S1**. Details of the literature data used in the analysis

| Modification | Approach | Source | Reference | Sample | Modified sites |
|---|---|---|---|---|---|
| **$N^6$-methyladenosine** | $m^6A$-CLIP-seq | GEO: GSE86336 | Ke et al., 2017 | HeLa | ~40000 |
| | TNT-seq | GEO: GSE83561 | Louloupi et al., 2018 | HEK293 | 58311 |
| **Pseudouridine** | CeU-seq | GEO: GSE63655 | Li et al., 2015 | HEK293 | 2058 |
| **A-to-I editing** | Matched RNA-Seq and whole genome sequence analysis | DREAM database | Picardi et al., 2015 | Human tissues | 4668508 |

**Table S2.** Fisher exact test results for PQS enrichment analysis

| Data | Observed PQS count | Expected PQS count | P-value | 95% CI | OR |
|---|---|---|---|---|---|
| $m^6A$ HEK293 pre-mRNA | 23372 | 5461 | 2.2 $e^{-16}$ | 6.27–6.69 | 6.47 |
| $m^6A$ HeLa pre-mRNA | 7919 | 3740 | 2.2 $e^{-16}$ | 2.32–2.52 | 2.42 |
| $m^6A$ HeLa mRNA | 7838 | 4131 | 2.2 $e^{-16}$ | 1.99–2.17 | 2.07 |
| Ψ HEK293 pre-mRNA | 308 | 195 | 8.836 $e^{-08}$ | 1.38–2.05 | 1.68 |
| Transcriptome A-to-I | 1004026 | 420067 | 2.2 $e^{-16}$ | 2.79–2.81 | 2.8 |

CI: confidence interval; OR: odds ratio 6.47.

**Table S3.** Two G-tetrad PQS loop nucleotide composition for those in $m^6A$-enriched regions

| | **Rank Order from Most to Least Common** | | | | | | | | | |
|---|---|---|---|---|---|---|---|---|---|---|
| | **1** | **2** | **3** | **4** | **5** | **6** | **7** | **8** | **9** | **10** |
| **LOOP 1 Identity** | **A** | **U** | **GA** | **G** | **AA** | **CU** | **CA** | **C** | **GU** | **AG** |
| **Count** | 1044 | 518 | 502 | 474 | 237 | 229 | 228 | 197 | 159 | 142 |
| **LOOP 2 Identity** | **A** | **U** | **G** | **GA** | **C** | **CUGA** | **AA** | **CA** | **CU** | **AG** |
| **Count** | 936 | 657 | 384 | 250 | 239 | 230 | 209 | 203 | 175 | 169 |
| **LOOP 3 Identity** | **A** | **U** | **G** | **CA** | **GA** | **AA** | **C** | **CU** | **AG** | **AU** |
| **Count** | 999 | 691 | 459 | 379 | 332 | 249 | 246 | 201 | 156 | 146 |

*Data corresponds to HEK293 pre-mRNA $m^6A$ profile reported by Louloupi, et al.[5]

**Table S4**. Frequency analysis of loop sequences in three or more G-tetrad PQSs in intronic m$^6$A enriched sites.*

| | 1 | 2 | 3 | 4 | 5 | 6 | 7 | 8 | 9 | 10 |
|---|---|---|---|---|---|---|---|---|---|---|
| **Loop1** | **A** | **CU** | **U** | **G** | **AA** | **CGU** | **CA** | **GA** | **AAG** | **UG** |
| **n** | 24 | 20 | 18 | 13 | 13 | 11 | 11 | 11 | 11 | 8 |
| **Loop2** | **A** | **U** | **CA** | **GU** | **CU** | **CCA** | **AG** | **C** | **GA** | **AU** |
| **n** | 30 | 19 | 15 | 13 | 13 | 39 | 9 | 8 | 7 | 7 |
| **Loop3** | **A** | **U** | **GA** | **AA** | **GU** | **CA** | **AGGA** | **AG** | **G** | **CU** |
| **n** | 26 | 18 | 18 | 16 | 12 | 11 | 11 | 10 | 9 | 9 |

*Data corresponds to HEK293 pre-mRNA m$^6$A profile reported by Louloupi, et al.[5]

**Figure S1**. Frequency of two and three or more G-tetrad PQSs in mature mRNA sequenced for $m^6A$, Ψ, and A-to-I editing.

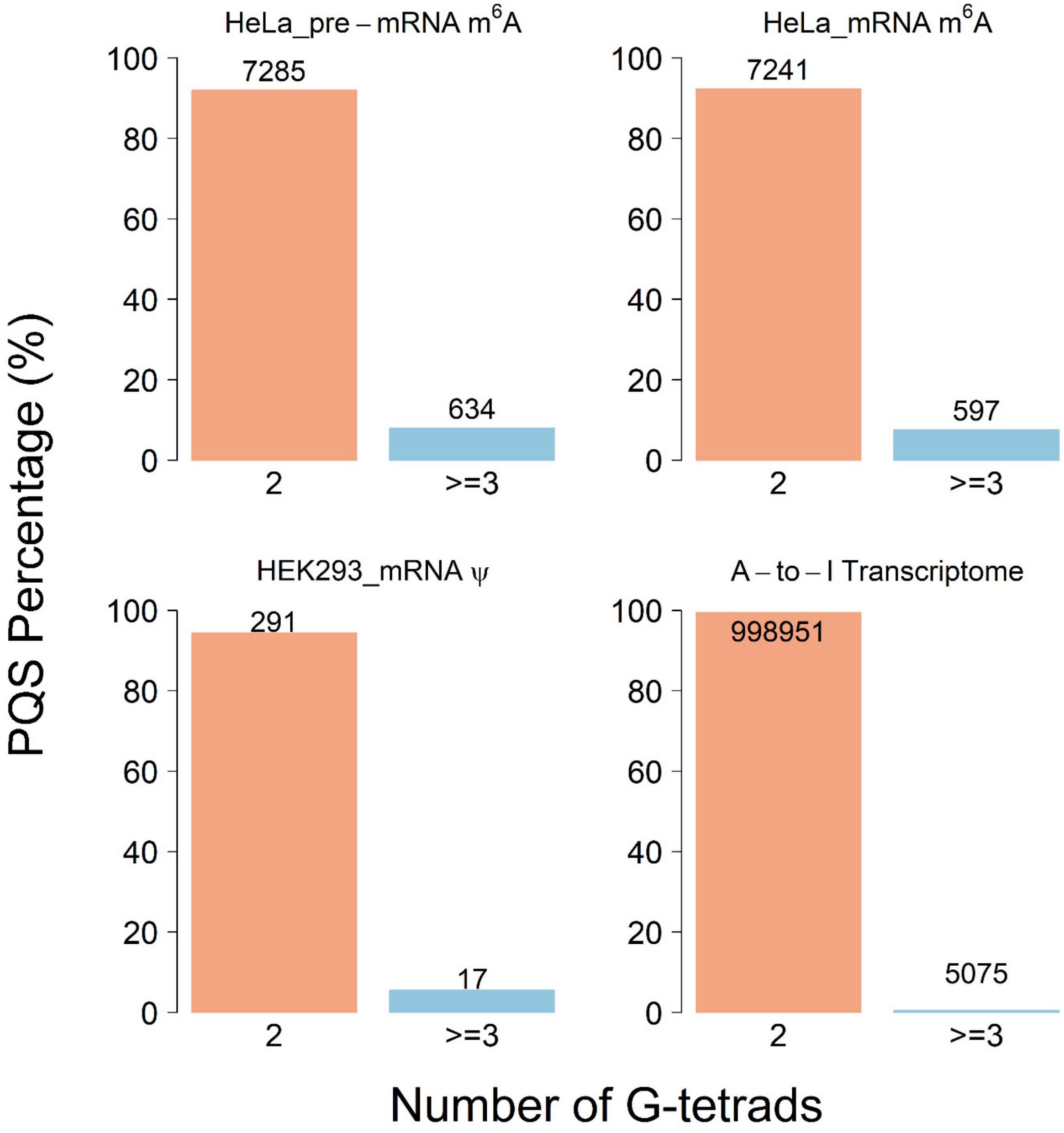
